# Low-coverage whole genome sequencing for a highly selective cohort of severe COVID-19 patients

**DOI:** 10.1101/2024.01.28.577610

**Authors:** Renato Santos, Víctor Moreno-Torres, Ilduara Pintos, Octavio Corral, Carmen de Mendoza, Vicente Soriano, Manuel Corpas

## Abstract

Despite advances in identifying genetic markers associated to severe COVID-19, the full genetic characterisation of the disease remains elusive. This study explores the use of imputation in low-coverage whole genome sequencing for a severe COVID-19 patient cohort. We generated a dataset of 79 imputed variant call format files using the GLIMPSE1 tool, each containing an average of 9.5 million single nucleotide variants. Validation revealed a high imputation accuracy (squared Pearson correlation ≈0.97) across sequencing platforms, showing GLIMPSE1’s ability to confidently impute variants with minor allele frequencies as low as 2% in Spanish ancestry individuals. We conducted a comprehensive analysis of the patient cohort, examining hospitalisation and intensive care utilisation, sex and age-based differences, and clinical phenotypes using a standardised set of medical terms developed to characterise severe COVID-19 symptoms. The methods and findings presented here may be leveraged in future genomic projects, providing vital insights for health challenges like COVID-19.

## Context

Coronavirus disease 2019 (COVID-19) is caused by the severe acute respiratory syndrome coronavirus 2 (SARS-CoV-2), which first appeared by the end of 2019 in Wuhan, China [1]. The clinical presentation of COVID-19 can be very heterogeneous, ranging from asymptomatic infection to severe forms with pneumonia, multiple organ complications, and sepsis [2]. Previous genome-wide association studies (GWAS) have collectively identified genetic risk factors at multiple loci across the human genome, including specific variants associated with COVID-19 severity and susceptibility to infection [3–5]. Additionally, certain patient characteristics, such as older age and male sex, as well as comorbidities, like obesity and cancer, have been shown to contribute to severe outcomes in COVID-19 patients [6]. This earlier body of work has paved the way for exciting new opportunities to explore the determinants of COVID-19 severity [7], particularly due to its potential applications in risk prediction, preventive medicine, and patient management.

Traditionally, the genotyping process has relied on array technologies as the standard, both at the broader GWAS level and the more specific genetic scoring and genetic diagnostics levels [8]. This reliance is primarily due to very low costs and fast turnaround times, which made microarrays valuable high-throughput tools capable of generating affordable genomic data. However, arrays are limited by their experimental design, leading to biases in the data generated. In particular, the prior selection of genetic markers and probes creates an ascertainment bias, resulting in the overrepresentation of intensively researched populations that are more likely to be involved in array development [9]. This contrasts with high-coverage whole genome sequencing (WGS), which, since its inception, has promised the ability to probe variation across the entire human genome, free from the ascertainment bias characteristic of microarrays. This, in fact, has led to its adoption by large scale population-level projects [10].

Despite significant cost reductions over time [11], WGS at the clinically accepted standard of 30X coverage [12] remains too expensive for many projects, especially those involving large sample cohorts such as those required for GWAS. However, recent studies have demonstrated that sequencing larger numbers of individuals at lower coverages, prioritising cost and haplotype diversity over sequencing depth, can actually–yield more allelic information at the cohort and population levels [13]. Consequently, low-coverage WGS (lcWGS) has emerged as a cost-efficient alternative to high-coverage WGS, surpassing microarrays in the discovery of common and low-frequency variation [14, 15], particularly in underrepresented populations [16].

Akin to microarrays, lcWGS data can also be imputed using reference panels to enhance resolution and statistical power while maintaining low sequencing and data processing costs. The fundamental principle that underlies genotype imputation algorithms is identity-by-descent (IBD), wherein two seemingly unrelated individuals may share segments shorter than 10 centimorgans (cM) inherited from a distant common ancestor [17]. Consequently, genotype imputation algorithms compare the sparsely distributed haplotypes present in the lcWGS data with the haplotypes in the high-coverage reference panel to infer genotype likelihoods in the regions not covered by sequencing [18].

Previous imputation methods for lcWGS data exhibited significant drawbacks that undermined their competitiveness. Some incurred higher costs and longer running times when using large reference panels due to computational complexity [19], and others used more efficient approximations, resulting in lower imputation accuracy [20, 21]. To address these challenges, we utilised the GLIMPSE1 algorithm [16], a less resource-intensive tool that produces more accurate imputed data than its predecessors, to generate a VCF dataset containing 79 imputed lcWGS samples, which we are releasing with this manuscript.

Although genotype imputation in lcWGS datasets shows promise, its use is still in the early stages. With continuous advancement in sequencing technologies, we expect that imputation methods will play an increasingly crucial role in unravelling the complexities of the human genome and accelerating discoveries in precision medicine and personalised healthcare.

## Dataset description

We generated a dataset consisting of 79 VCF files, and respective FASTQ and CRAM files, using the GLIMPSE1 imputation algorithm [16]. We leveraged the 1000 Genomes Project Phase 3 dataset [22] as the reference panel of haplotypes. In total this dataset is composed of approximately 325 GB of FASTQ data, 156 GB of CRAM data, and 6 GB of VCF data.

Our samples were specifically derived from sequenced DNA from a highly selective cohort of patients, comprised of mostly Iberian Populations in Spain (IBS) individuals but also containing some individuals from other genetic backgrounds. All patients presented with severe COVID-19 symptoms during the initial wave of the SARS-CoV-2 pandemic in Madrid, Spain.

On average, each VCF file in this rich dataset contains 9.49 million high-confidence single nucleotide variants [95%CI: 9.37 million - 9.61 million] (**Figure 1**). To facilitate access to researchers interested in further studying this data, it has been made available for reuse in the European Genome-phenome Archive [23], under the accession number EGAS00001007573.

**Figure 1.**
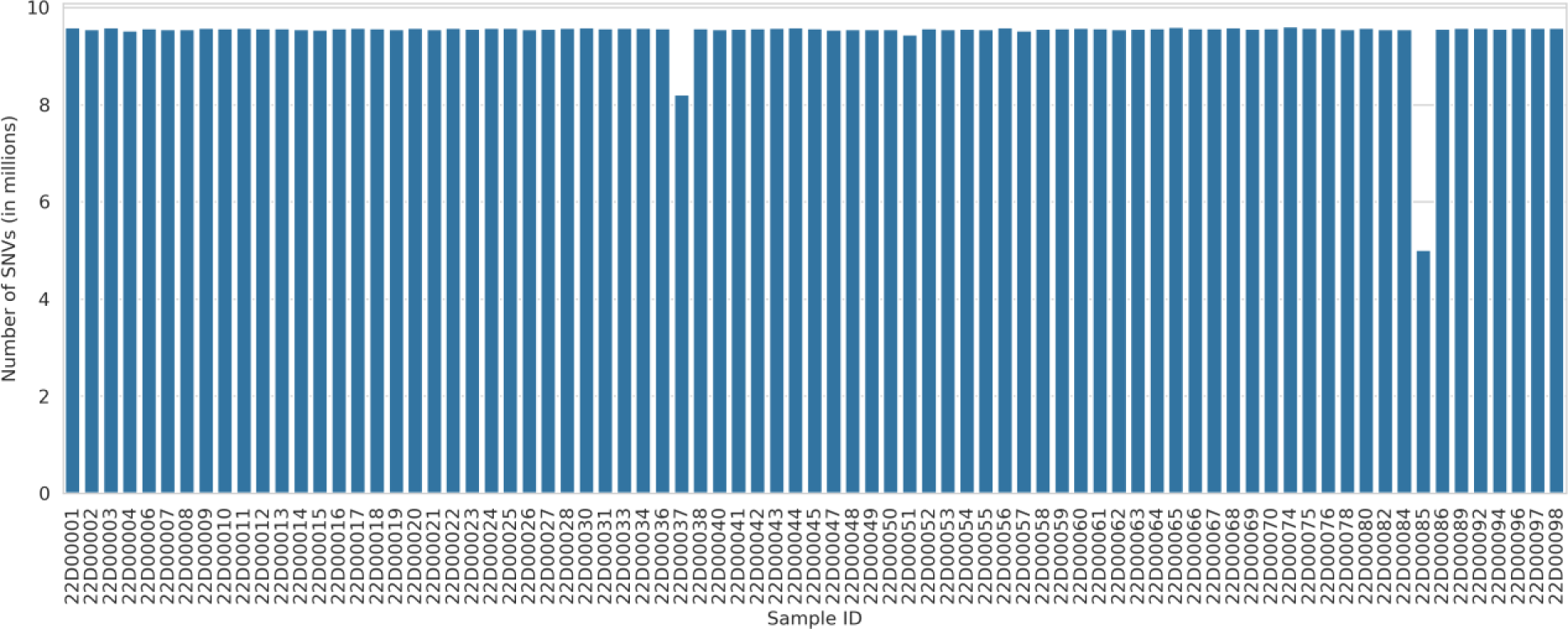
Number of high-confidence single nucleotide variants (SNVs) for the 79 VCF files in the severe COVID-19 dataset. The x-axis represents the sample IDs in the dataset, while the y-axis denotes the total counts of SNVs for each sample in millions (1×10^6^).

## Patient cohort characterisation

### Sampling strategy

The 79 genomic samples analysed in this study constitute a subset of a larger cohort of individuals whose exomes were initially sequenced and analysed as part of a comprehensive investigation into genetic determinants for COVID-19 severity [24]. The selection of this subset was based on the quality assessment of DNA samples suitable for PCR-free library preparation for lcWGS. All individuals were patients hospitalised between March and June 2020, during the first wave of the SARS-CoV-2 pandemic in Spain, at a tertiary referral hospital in Madrid, and confirmed to be infected with SARS-CoV-2. We aimed to select patients with the following clinical profile: (1) younger than 60 years old; (2) experienced fever and respiratory symptoms for more than three days; (3) blood oxygen saturation level below 93%; (4) bilateral pneumonia on imaging tests; and (5) no comorbidities, such as diabetes, obesity, or immunosuppressive conditions. At the time the study population was recruited, no vaccines had been developed yet.

We examined the dataset focusing on three main aspects: firstly, a general characterisation of the patients by age, sex, and ethnicity; secondly, hospital stays and time spent in the intensive care unit (ICU); and finally, the distinct clinical phenotypes presented by the patients (respiratory, thromboembolic, cardiovascular, etc.). Further details on the patients’ demographic information and clinical history can be found in **Supplementary File 1** [25].

### Demographic characterisation

Through our analysis, we aimed to create a comprehensive demographic profile of our cohort of severe COVID-19 patients. The age distribution in the cohort (**Figure 2A**) is characterised by a distinct right skew, with a higher prevalence of individuals falling within the 45-64 age bracket and particularly concentrated around 55-59 years, which aligns with our current knowledge of the correlations between older age and severe COVID-19 outcomes [6]. Yet, the lower tail-end of the distribution also underscores the fact that severe COVID-19 is not strictly age related and young individuals may also suffer from severe manifestations.

The sex distribution (**Figure 2B**) shows a higher frequency of male patients relative to females. This finding concurs with previous research indicating that men are at a higher risk of developing severe COVID-19 [6]. Investigating the age distribution in relation to sex (**Figure 2C**), indicates that both males and females have a similar median age of 55 and 53 years, respectively, although the male age distribution exhibits a broader range and higher variability, suggestive of a greater scope of age-related COVID-19 risk among men.

Lastly, studying the patients’ country of origin (**Figure 2D**) reveals that most of the patients in our cohort originate from Spain and Latin American countries. This geographical distribution is mostly a reflection of the demographic makeup of Madrid, Spain, where sample collection occurred. To expand our analysis beyond demographics and understand the genetic makeup of our cohort, we also performed a Principal Component Analysis (PCA) of our 79 samples, after imputation and variant filtering, against the backdrops of the 1000 Genomes Project [22] global superpopulations and IBS population.

The global PCA plots (**Figure 3A**) show that most samples cluster within the European (EUR) group, mirroring the fact that a significant proportion of our cohort hails from Spain. Additionally, a subset of patients is found within or near the Admixed American (AMR) and South Asian (SAS) clusters, reflecting the Latin American patients in our cohort, and the mixed ancestry common in Latin American populations. A few patients also cluster within the African (AFR) group, likely representing the African ancestry in our cohort from the patients originating from Cape Verde and Morocco.

In the IBS-specific PCA plots (**Figure 3B**), most of the severe COVID-19 patients form a distinct cluster close to the 1000 Genomes IBS population, indicating a shared genetic background with this group, representing the individuals with IBS ancestry. However, it is worth noting that subtle regional genetic variations within the Iberian population could contribute to the observed dispersion within this shared genetic background, particularly along the third principal component. The figure also shows a dispersion of patients alongside the left side of the plots, which represent the individuals with ancestries other than IBS. This highlights the genetic diversity in our cohort, contributed by the inclusion of patients from Spain and various other nations.

**Figure 2.**
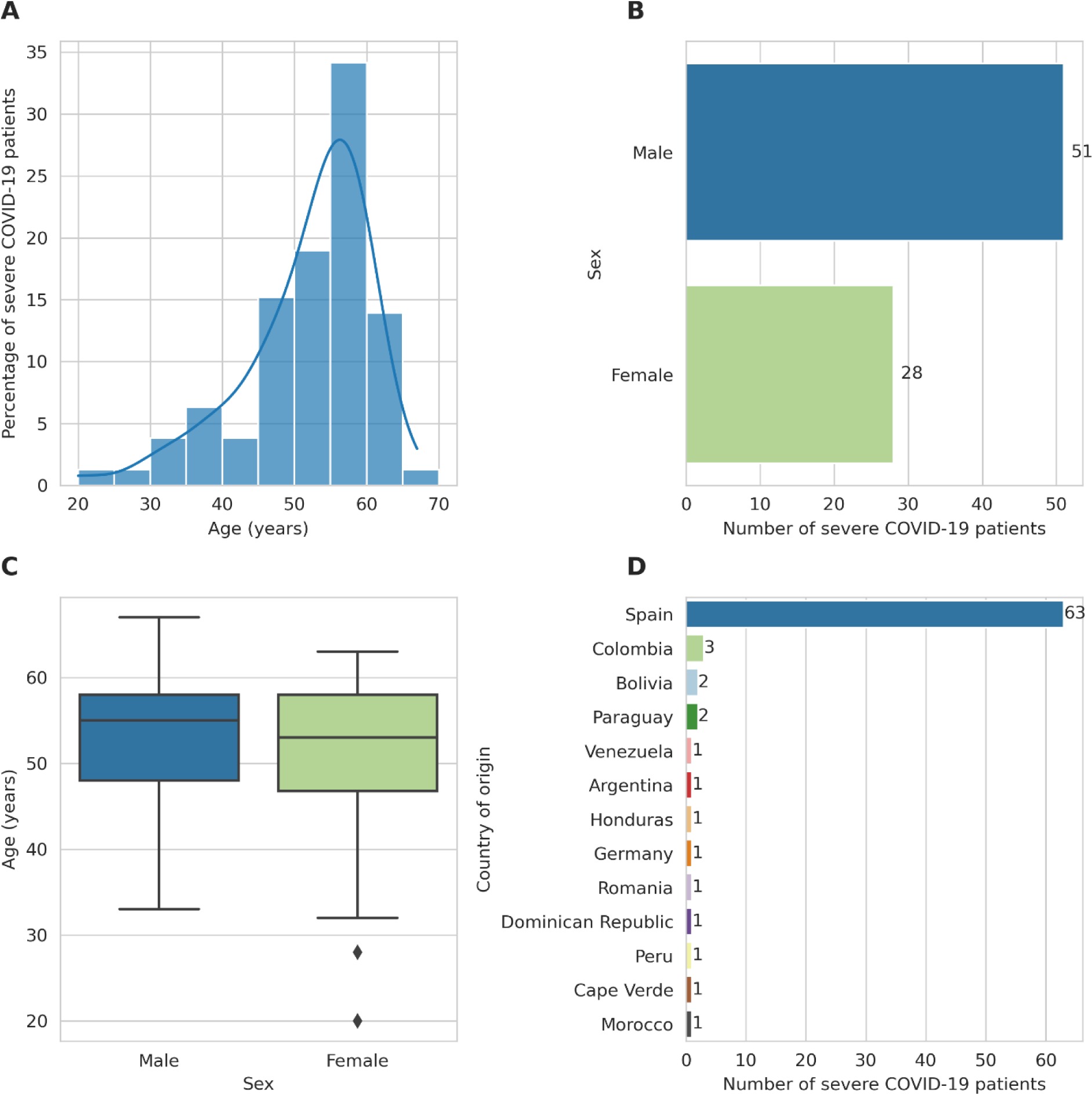
Demographic and geographic characterisation of the severe COVID-19 patient cohort. **A** Distribution of severe COVID-19 patients’ ages in our cohort. Each bar signifies an age bracket comprising 5-year increments, with its height denoting the proportion of individuals within that age range. The plot is overlaid with a Kernel Density Estimation (KDE) curve, which provides a smoothed estimation of the age distribution. **B** Patients’ stratification by sex. Each bar represents one sex, with its length indicating the number of patients of that sex. **C** Distribution of patients’ ages by sex. The boxplot presents the age distribution for each sex. Each box represents the interquartile range (IQR) of ages for either males or females, with the dividing line representing the median age. The diamonds represent outliers. **D** Distribution of patients by country of origin. Each bar corresponds to a country, and its length indicates the number of patients from that country.

**Figure 3.**
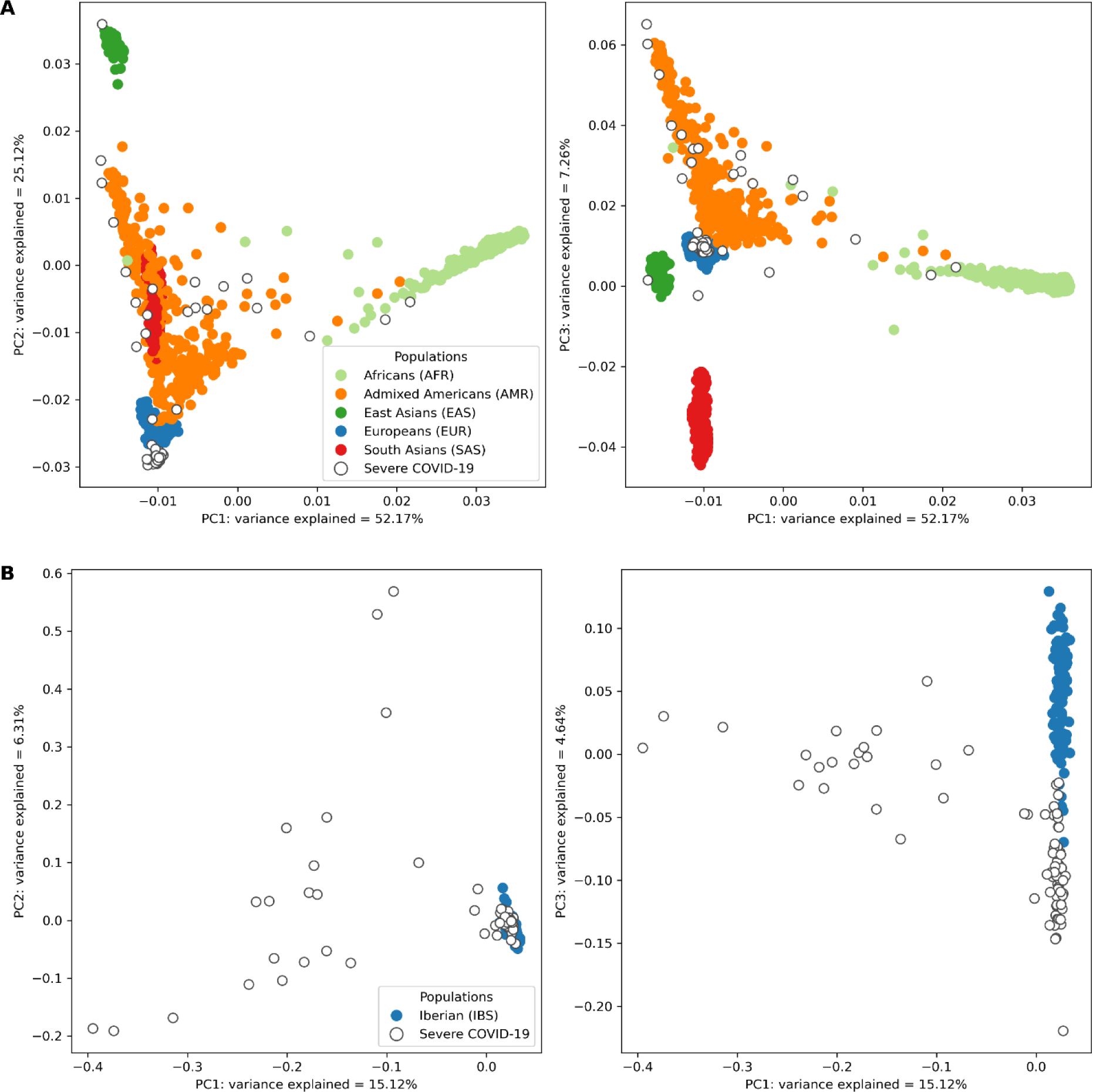
Principal component analysis of genetic variation in the severe COVID-19 patient cohort against the 1000 Genomes Project global superpopulations and IBS (Iberian Populations in Spain) population. **A** Projection of imputed low-coverage whole-genome sequencing (lcWGS) data from severe COVID-19 patients against the backdrop of global superpopulations from the 1000 Genomes Project. Each point represents an individual, colour-coded according to their superpopulation. Severe COVID-19 patients are distinguished by points with a white fill and coloured border. The x-axis and y-axis on the two subplots represent the first and second, and first and third principal components, respectively, with the percentage of variance explained by each component indicated in the axis label. **B** Focused view of the genetic variation within the Iberian (IBS) population and the severe COVID-19 patients. Individuals from the IBS population are represented by solid-coloured points, while those with severe COVID-19 are represented by points with a white fill and coloured border. The x-axis and y-axis on the two subplots represent the first and second, and first and third principal components, respectively, with the percentage of variance explained by each component indicated in the axis label.

### Hospital stays

Examining the patients’ hospital medical records provides valuable insights about the hospitalisation experience of individuals with severe COVID-19. By examining these trends, we can gain a better understanding of potential sex and age-based differences in the duration of hospitalisation and the level of care required.

Firstly, we analysed the distribution of hospitalisation days in our patient cohort (**Figure 4A**). The distribution is notably skewed to right, with most patients requiring relatively short hospital stays between 1 and 34 days. However, the distribution’s right tail shows that a subset of patients experienced significantly longer stays, up to 202 days, which could be attributed to cases of COVID-19 with increased severity, thus requiring additional medical attention.

Furthermore, an evaluation of distribution of hospital stays by sex (**Figure 4B**) reveals that the median duration of hospital stays was similar for both sexes. Nevertheless, the distribution for male patients exhibits greater variability, accentuated by the presence of some outliers who spent an unusually high number of days in the hospital, which represent severe or complex cases that required a significantly longer time for recovery and medical management. This could mean that the severe disease progression and recovery time in males is less consistent than in females, possibly due to a wider range of severity in clinical presentations among male patients.

In addition, we investigated the use of the intensive care unit (ICU). Approximately 25% of the cohort was admitted to the ICU during their hospitalisation (**Figure 4C**), indicating that, despite the severity of their COVID-19 symptoms, most patients were managed without the need for intensive care. However, a much larger proportion of males necessitated ICU admission than females (**Figure 4D**). This, again, reflects findings from numerous studies that have identified male sex as a risk factor for severe COVID-19 outcomes [6].

We further stratified the ICU data by patient age (**Figure 4E**), showing that the majority of patients who were admitted to the ICU were between 45 and 70 years old, which underscores the heightened risk of severe outcomes in these age groups. Finally, we investigated the duration of ICU stays among those who required such care (**Figure 4F**). Mirroring the hospital stay duration for the overall cohort, most patients admitted to the ICU spent between 1 and 35 days there. However, a considerable subset of patients experienced significantly longer ICU stays, thus representing a broad spectrum of disease severity and recovery rates within this critical care cohort.

**Figure 4.**
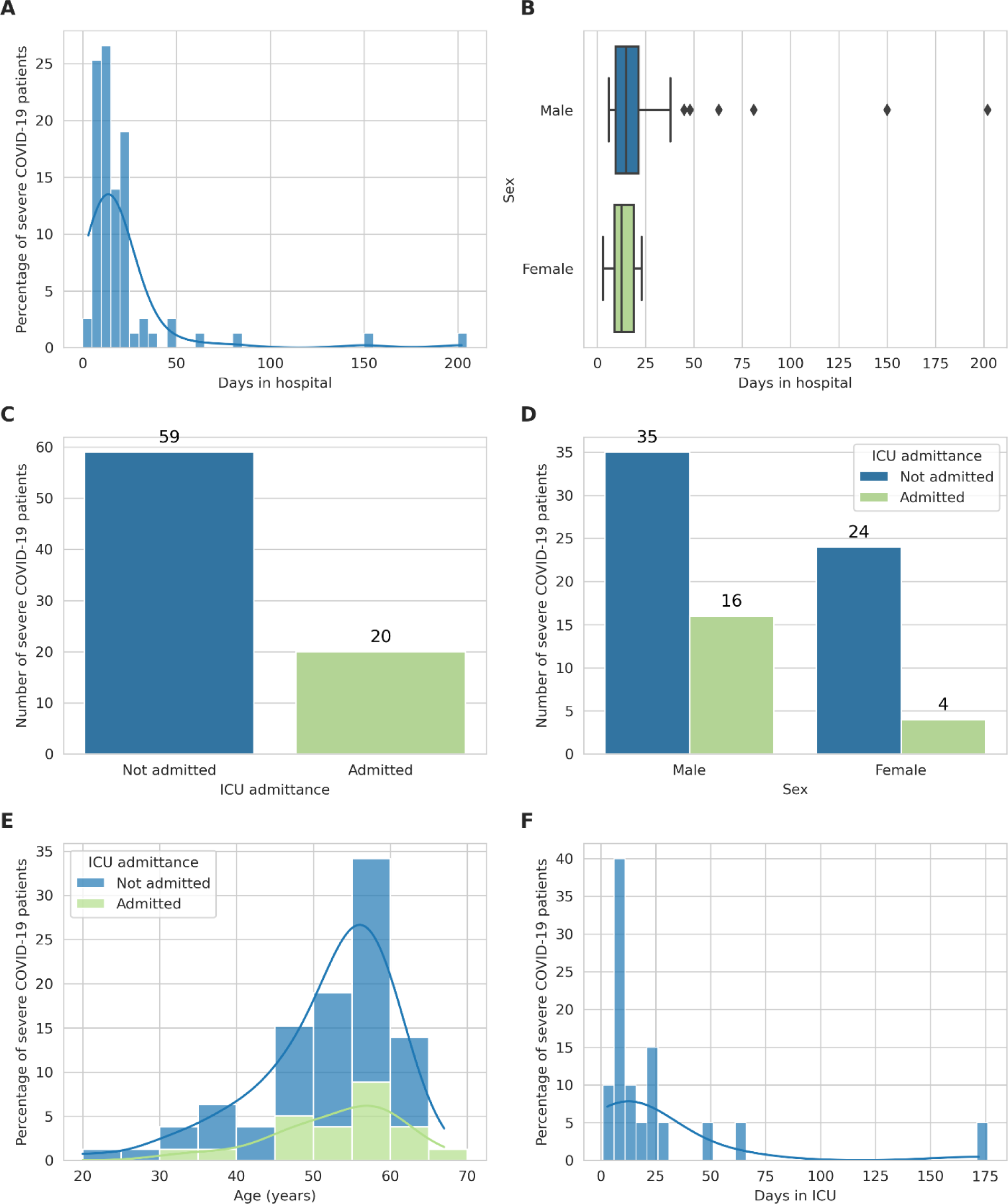
Analysis of hospital stays among the severe COVID-19 patient cohort. **A** Distribution of hospital stay durations in our cohort. Each bar corresponds to an interval of hospital stay durations of 5 days, with its height indicating the proportion of patients with a stay duration within that duration interval. The plot is overlaid with a Kernel Density Estimation (KDE) curve, that provides a smoothed estimate of the duration distribution. **B** Stratification of hospital stay durations by sex. This boxplot presents the distribution of hospital stays for each sex. Each box represents the interquartile range (IQR) of the duration of hospital stays for one sex, with the line inside the box marking the median duration. The diamonds represent outliers. **C** Distribution of patients admitted to the Intensive Care Unit (ICU). Each bar corresponds to either patients admitted to the ICU (green) or patients not admitted to the ICU (blue), with its height indicating the number of patients in that category. **D** Distribution of patients admitted to the ICU by sex. Each pair of bars corresponds to one sex, with their height indicating the proportion of patients of that sex who were admitted to the ICU. Each bar corresponds to either patients admitted to the ICU (green) or patients not admitted to the ICU (blue), with its height indicating the number of patients in that category. **E** Distribution of ages of patients admitted to the ICU. Each bar corresponds to an age group of 5 years, with its height indicating the proportion of patients in that age group. The plot is overlaid with a KDE curve, which provides a smoothed estimate of the age distribution. **F** Distribution of ICU stay durations among patients admitted to the ICU. Each bar corresponds to an interval of ICU stay durations of 5 days, with its height indicating the number of patients within that duration interval. The plot is overlaid with a KDE curve, that provides a smoothed estimate of the duration distribution. Only patients who were admitted to the ICU are represented in this plot.

### Clinical phenotypes

An analysis of the phenotypes of the severe COVID-19 patient cohort reveals valuable insights into the most common phenotypes associated with severe forms of the disease and their frequency and relationships. While established COVID-19 phenotype ontologies were readily available [26, 27], they lacked the level of granularity we required to comprehensively characterise the clinical phenotypes of our cohort. Therefore, we devised a specialised set of standardised terminology comprising 28 medical terms that were organised into 4 primary clinical categories: pulmonary, extra-pulmonary, coagulation, and systemic phenotypes. Subsequently, we evaluated each patient’s record for the presence of these terms.

**Table 1** provides a detailed breakdown of the number of patients associated with each specific phenotype, within the four major clinical categories. We found that the Pulmonary category, which includes pneumonia, ARDS (acute respiratory distress syndrome), and a combination of ARDS and admission to the ICU, was the most prevalent among our cohort. Indeed, pneumonia alone was identified in 78 patients. The Extra-Pulmonary category covers a broad range of clinical symptoms and conditions, with liver hepatitis and gastrointestinal diarrhoea being the most common, observed in 10 patients each. The Coagulation category focused on thrombotic events and related conditions. Pulmonary embolism and deep venous thrombosis, each identified in 5 patients, were most prevalent. Finally, the Systemic category included conditions that affect the patient’s overall health and wellbeing, such as persistent fever and symptoms like fatigue and headache. Persistent fever was the most common Systemic phenotype, observed in 33 patients.

To further investigate the relationships between the phenotypes in our patient cohort and to determine whether any of them were likely to co-occur, we performed a Spearman correlation analysis using the function *corr(method=“spearman”)* from the seaborn package for Python [28], and visualised the results in a heatmap (**Figure 5**). These correlations suggest that patients who presented with one of these phenotypes were more likely to present others, pointing to possible common underlying pathways or simultaneous occurrence in severe disease presentation.

The plot shows that most phenotypes are not strongly correlated, hence, the presence of one phenotype does not necessarily predict the presence of another. This could be indicative of the diverse clinical manifestations of severe COVID-19, with different phenotypes appearing independently in different patients. However, there are several pairs of phenotypes exhibiting higher degrees of correlations. This is particularly evident in neurological conditions, such as the correlations between psychiatric disorders, encephalopathies and polyneuropathies, which appear to be correlated to a relatively high degree. In addition, moderate correlations are shown between the former three neurological phenotypical categories and exanthema, myopathies, and bone marrow abnormalities. Finally, some moderate correlations occurred between the ARDS & ICU phenotype and a few other phenotypes, pointing at the additional occurrence of various phenotypes of COVID-19 severity in patients admitted to the ICU with ARDS.

This exploratory analysis highlights the diverse ways in which severe COVID-19 can present, and the importance of comprehensive and nuanced clinical phenotyping in improving our understanding and management of the disease.

**Table 1.**
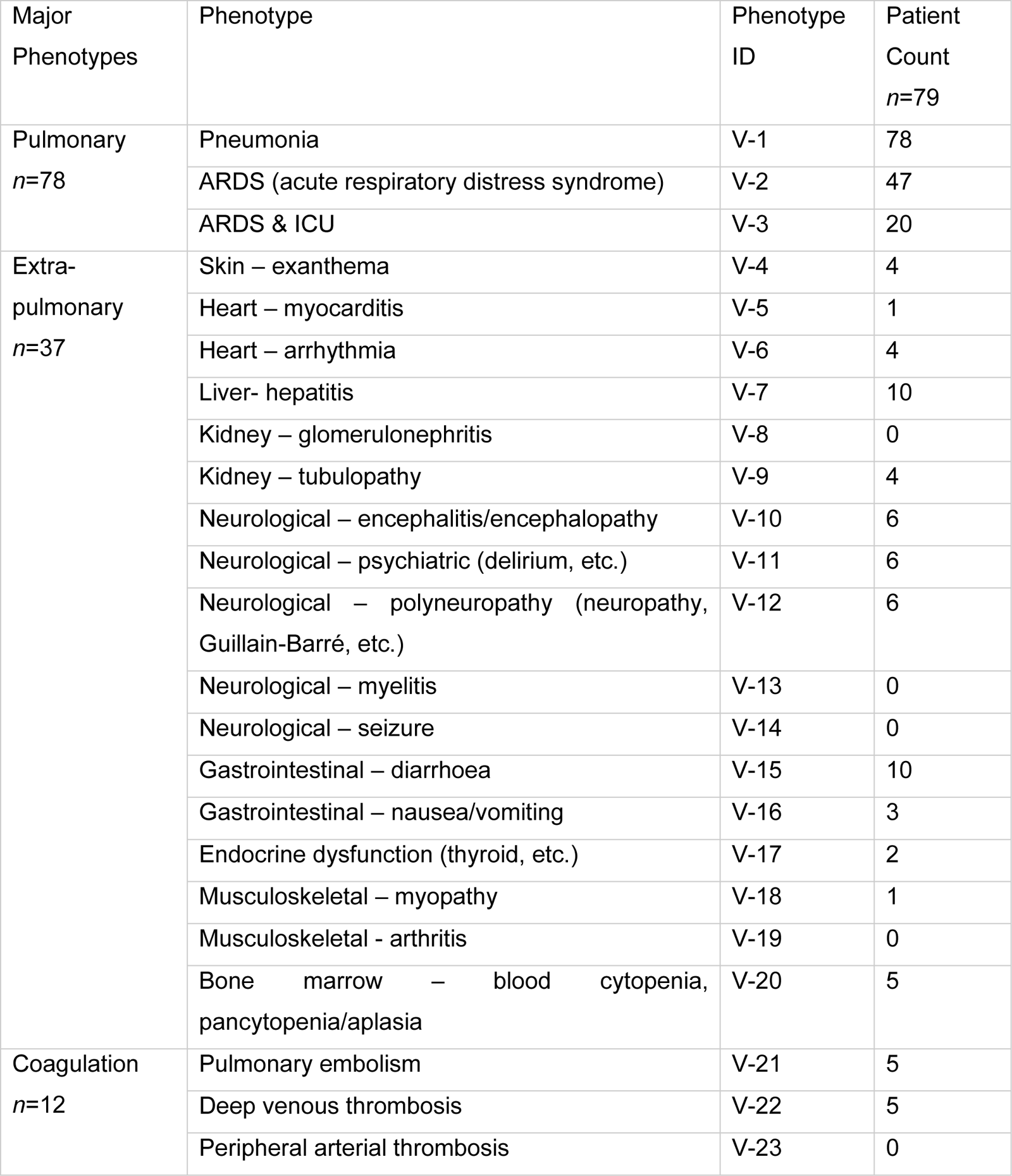
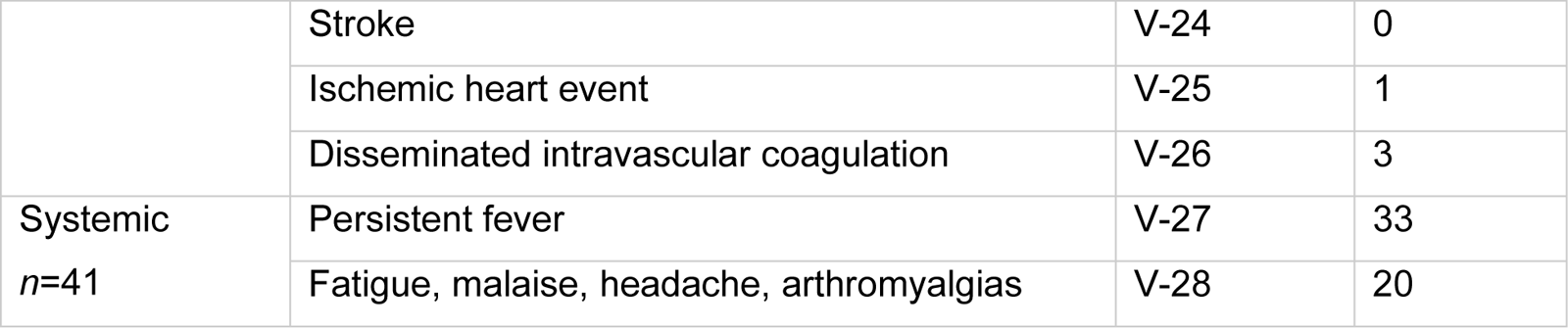
Frequency of severe COVID-19 phenotypes in the patient cohort. The table presents the distribution of our 28 severe COVID-19 specific phenotypes organised into four major clinical categories: Pulmonary, Extra-pulmonary, Coagulation, and Systemic, as observed in our severe COVID-19 patient cohort. For each category, the total number of unique patients with at least one phenotype in the category is indicated. Each phenotype is listed with a unique Phenotype ID (V-1 to V-28) and the number of patients who were identified with that phenotype.

**Figure 5.**
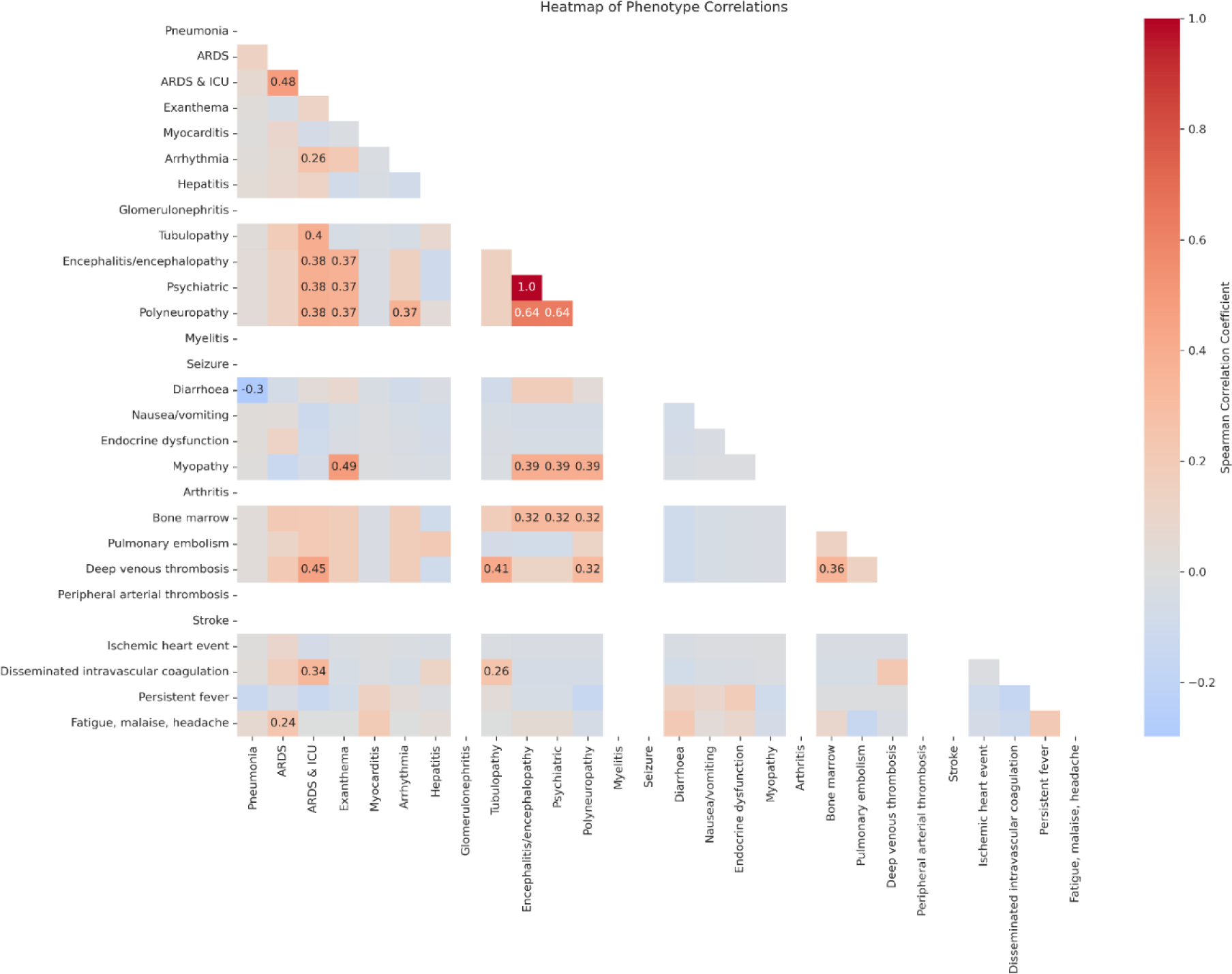
Heatmap of phenotype correlations in the severe COVID-19 patient cohort. The plot illustrates the Spearman correlations between our 28 severe COVID-19 specific phenotypes. Each square in the heatmap represents the correlation between two phenotypes, with the colour of the square indicating the strength of the correlation, and the number inside each square represents the correlation coefficient. Only statistically significant (p < 0.05) correlation coefficients are shown. Darker colours represent stronger positive or negative correlations, with red representing positive correlations and blue representing negative correlations. These indicate that individuals displaying one phenotype are more or less likely to exhibit the other phenotype as well, suggesting potential underlying mechanisms for the progression of severe COVID-19.

## Methods

### DNA extraction and library preparation

We collected blood samples for each patient in 10 mL EDTA tubes. We then centrifuged the tubes at 3000 rpm for 10 minutes to isolate the buffy coat, which we subsequently froze at -20°C until further use. Afterward, we used the Maxwell RSC Buffy Coat DNA Kit (AS1540, Promega UK, Southampton, United Kingdom) to isolate genomic DNA from frozen buffy coat samples. We assessed the concentration of the genomic DNA using spectrometric analysis. We then fragmented the DNA using a Covaris E220 focused-ultrasonicator (Covaris Ltd., Brighton, United Kingdom) to generate 350 bp length DNA fragments. The following parameters were used for the fragmentation process: *6 cycles*, *PIP 75*, *Cycles/Burst 1000*, *Duty Factor 20%*, *Duration 20s*. After that, we prepared DNA libraries using the MGIEasy PCR-Free DNA Library Prep Set (1000013453; MGI Tech Co. Ltd, Shenzhen, China). The concentration of the DNA libraries was assessed using a Qubit 3.0 fluorometer (Life Technologies). Finally, we sequenced the libraries on an MGI DNBSEQ-G400 sequencer (RRID:SCR_017980; MGI Tech Co. Ltd), with a target sequencing depth of 1X.

### Sequencing quality control and preprocessing

We performed quality control and preprocessing of the resulting FASTQ files using the nf-core [29] Sarek pipeline v3.1.2 [30–39]. The following parameters were applied during the pipeline execution: *nextflow run nf-core/sarek -r 3.1.2 -profile docker --input samplesheet.csv –outdir /mnt/e/Sarek/out/ --trim_fastq --igenomes_base /mnt/e/Sarek/references --genome GATK.GRCh37 --skip_tools strelka --seq_platform ‘MGI’*. We used the recalibrated base quality scores CRAM files produced by the Sarek pipeline as the input for the subsequent step.

### Imputation using GLIMPSE

We calculated genotype likelihoods using bcftools mpileup v1.16 (RRID:SCR_005227) [39], with the parameters *-I -E -a ‘FORMAT/DP’*, followed by genotype calling using bcftools call, with parameters *--ploidy GRCh37 -S ploidy.txt -Aim -C alleles*. The file ‘ploidy.txt’ contained information about the sex of each sample, which was necessary to correctly generate genotype calls for chromosome X in males [16].

After that, we imputed the low-coverage genomes with GLIMPSE v1.1.1 [16]. Firstly, we split each chromosome into 2 Mb chunks, with 200 kb buffer regions on each side of a chunk, by using GLIMPSE_chunk with the parameters *--window-size 2000000 --buffer-size 200000*. Secondly, we used GLIMPSE_phase to perform imputation of each chunk, with the tool’s default iteration parameters. GLIMPSE_phase imputation was multithreaded with GNU Parallel v20230522 [40].

We used the 1000 Genomes Phase 3 dataset [22] as the reference panel, since it has been shown that the inclusion of diversity in reference panels improves the quality of imputation by reducing missing genotype calls [41, 42]. Finally, we used GLIMPSE_ligate to join the imputed chunks together into entire chromosomes, followed by bcftools concat to merge all chromosomes into a single VCF file, containing chromosomes 1 to 22 and X, for each sample.

### Post-imputation filtering

Following imputation, we filtered the VCF files, to prioritise the most reliable genotype calls for further analysis, keeping only variants with minor allele frequency (MAF) above 2% and maximum genotype probability (GP) above 80%. We determined the MAF value through our validation process (see Data Validation) as the minimum threshold of acceptable imputation accuracy of *r*^2^ ≅ 0.9. The GP field represents the likelihood of each genotype being accurate and is expressed as a value between 0 and 1, with the sum of probabilities across all possible genotypes totalling 1 [43]. The chosen cutoff was determined as the best compromise between imputation accuracy and loss of information [43]. With this approach, we generated a dataset of imputed VCF files, from the DNA samples of our severe COVID-19 cohort.

### Principal component analysis

We performed a principal component analysis (PCA) to assess the genetic ancestries of our patient cohort, using PLINK v.1.90b6.21 (RRID: SCR_001757) [44]. To do so, firstly we normalised the variants from chromosomes 1 to 22 of the 1000 Genomes Project Phase 3 dataset [22] using bcftools [39]. We split multi-allelic calls and left-aligned indels against the reference genome using bcftools norm, with the parameters *-m-any --check-ref w*; followed by bcftools annotate with the parameters *-x ID -I +’%CHROM:%POS:%REF:%ALT’*, to normalise the naming of unset IDs; and bcftools norm to remove duplicate records using the parameters *-Ob --rm-dup both*. In addition, we filtered this global variant dataset to create a distinct subset containing variants exclusively from the 1000 Genomes Project IBS individuals. We used bcftools view, with the *-S* parameter, which includes only variants originating from the specified 1000 Genomes IBS sample IDs.

We then converted the two datasets, 1000 Genomes global and IBS, to binary PLINK format, using PLINK with the parameters *--keep-allele-order --vcf-idspace-to _ --const-fid --allow-extra-chr 0 --split-x b37 no-fail --make-bed*. Next, we used PLINK to identify population-specific markers by filtering the variants within both datasets based on MAF and variance inflation factor (VIF), using the parameters *--maf 0.10 --indep 50 5 1.5*. We then pruned the datasets to include only the variants that fulfil those thresholds, through *–extract*.

Subsequently, we merged the variants from chromosomes 1 to 22 of all 79 post-filtering imputed VCF files into a single BCF file using bcftools merge with the parameters *-r $(seq -s, 1 22) -- missing-to-ref*, followed by a normalisation process akin to that of the 1000 Genomes global and IBS datasets. Then, we converted the merged imputed dataset to binary PLINK format.

We used PLINK again to determine common variants between the merged imputed dataset and the populational markers in the 1000 Genomes global and IBS datasets, using *–bmerge*, followed by extraction of common variants with *–extract*. Finally, we merged the extracted variant datasets with *–bmerge*, and calculated 20 principal components using *–pca*. We plotted the first three principal components using the matplotlib package for Python [45].

## Data Validation

To validate our imputed dataset, we sought to quantify the accuracy of our imputation process and determine whether it is affected by the choice of short-read sequencing platforms. To do so, we obtained a healthy IBS genome (IBS001), independent from our patient cohort, sequenced at 40X coverage using an Illumina system, at Dante Labs (Cambridge, United Kingdom), and an MGI system, at BGI Tech Solutions Hongkong Co Ltd (Tai Po, Hong Kong). After quality control, alignment, and pre-processing, we obtained two CRAM files for the IBS001 individual: *IBS001_illumina* with an average depth per base of 39.34 and *IBS001_bgi* with an average depth per base of 41.64. The average depth per base was calculated using mosdepth v0.3.3 [36], with the *-x* parameter, against the GATK b37 reference genome.

We downsampled both versions of the IBS001 genome to 1X coverage using samtools v1.16 (RRID:SCR_002105) [39]. We calculated the subsampling fractions by dividing the target depth per base (1.00) by the respective average depth per base of each file. Therefore, the genome sequenced on the Illumina platform was downsampled using the command *samtools view -s 0.025419420437214,* and the genome sequenced on the MGI platform was downsampled using *samtools view -s 0.0240153698366955*.

The two resulting low-coverage downsamples were used for the genotype calling, imputation, and post-imputation filtering steps as described in the *Methods* section. This process resulted in four VCF files: two pre-filtering files and two post-filtering files from the two sequencing platforms, Illumina and MGI. These four files served as the imputed genotypes for the validation process. Additionally, we also performed the genotype calling step on the original high-coverage files, with the VCF output used as the gold standard genotypes in the validation.

To measure imputation accuracy we used GLIMPSE_concordance [16] to calculate a squared Pearson correlation between the high-coverage and imputed dosages across chromosomes 1 to 22 and X, within several MAF bins (0-0.02%, 0.02-0.05%, 0.05-0.1%, 0.1-0.2%, 0.2-0.5%, 0.5-1%, 1-2%, 2-5%, 5-10%, 10-15%, 15-20%, 20-30%, 30-40%, and 40-50%), and a single aggregate squared Pearson correlation across all sites. We only used sites in the validation data with a minimum depth of 8 reads and minimum posterior probability of 0.9999. To do so, we used the parameters *--minDP 8 --minPROB 0.9999 --bins 0.00000 0.0002 0.0005 0.001 0.002 0.005 0.01 0.02 0.05 0.1 0.15 0.2 0.3 0.4 0.5*.

We conducted this concordance assessment, comparing the imputed pre-filtering IBS001 genomes against the high-coverage validation IBS001 genomes, within sequencing platforms (BGI 1X vs BGI 40X and Illumina 1X vs Illumina 40X) and across sequencing platforms (BGI 1X vs Illumina 40X and Illumina 1X vs BGI 40X), to identify any quantifiable differences in imputation quality arising from the use of different short-read sequencing platforms. Across all four platform comparisons (**Figure 6A**), GLIMPSE accurately imputed variants in the 2-5%, 5-10%, 10-15%, 15-20%, 20-30%, 30-40%, and 40-50% MAF bins, represented by an r^2^ correlation equal to or higher than 0.90. However, as anticipated, imputation accuracy steadily decreased for MAFs lower than 2%, likely due to the underrepresentation of rare variants in the 1000 Genomes reference panel [16]. Overall, this resulted in an aggregate r^2^ correlation of approximately 0.96 across all MAF bins for both IBS001 genomes (**Table 2**).

Moreover, we assessed the impact of the filtering process on the accuracy of the dataset. We repeated the concordance comparison using GLIMPSE_concordance, instead using the imputed post-filtering IBS001 genomes against the high-coverage validation IBS001 genomes. Due to the removal of low confidence sites during filtering, the r^2^ correlation in the 2-5%, 5-10%, 10-15%, 15-20%, 20-30%, 30-40%, and 40-50% MAF bins improved slightly (**Figure 6B**). In turn, this improved the aggregate r^2^ correlation to approximately 0.97 on all four platform comparisons we performed (**Table 2**).

In conclusion, the validation of our imputation and filtering process shows that GLIMPSE1, with the 1000 Genomes Project Phase 3 [22] as the reference panel, can be used to confidently impute variants with MAF up to approximately 2%.

**Figure 6.**
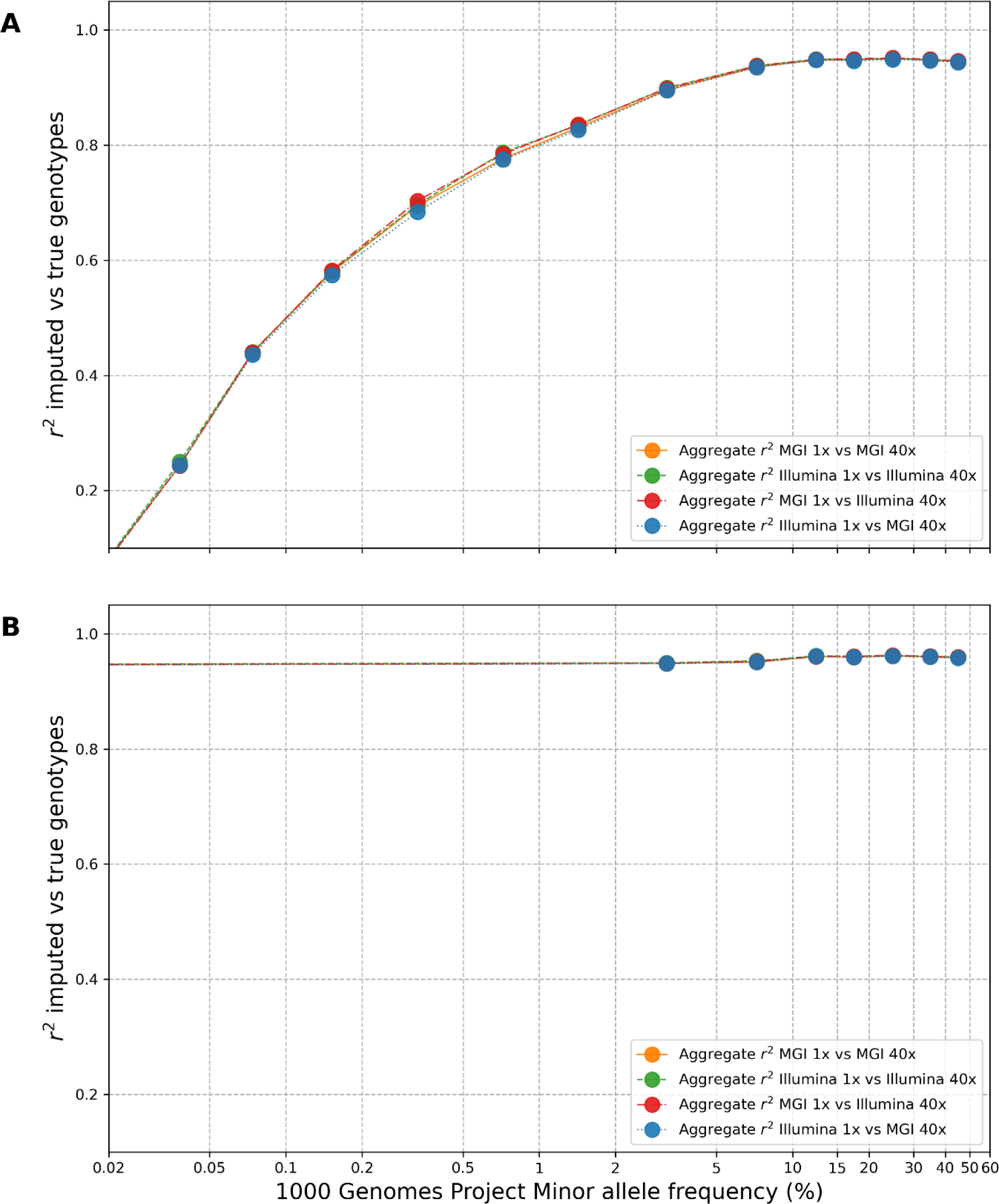
Assessment of GLIMPSE1 imputation concordance within different minor allele frequency (MAF) bins for the IBS001 validation genome. **A** Squared Pearson correlation (r^2^) between high-coverage and pre-filtering imputed dosages segregated into various MAF bins. The x-axis shows MAF bins, ranging from 0 to 50%, and the y-axis shows the squared Pearson correlation coefficient (r^2^). The analysis was performed for chromosomes 1 to 22 and X, within sequencing platforms (BGI 1X vs BGI 40X and Illumina 1X vs Illumina 40X) and across sequencing platforms (BGI 1X vs Illumina 40X and Illumina 1X vs BGI 40X). **B** Squared Pearson correlation (r^2^) between high-coverage and post-filtering imputed dosages segregated into various MAF bins.

**Table 2.** Aggregate GLIMPSE1 imputation concordance for the IBS001 validation genome. The table displays the aggregate r^2^ correlation results obtained from the concordance assessment of the imputed IBS001 genomes against the high-coverage validation IBS001 genomes. The analysis was performed for chromosomes 1 to 22 and X, within sequencing platforms (BGI 1X vs BGI 40X and Illumina 1X vs Illumina 40X) and across sequencing platforms (BGI 1X vs Illumina 40X and Illumina 1X vs BGI 40X). The table presents the aggregate r^2^ correlation values, indicating the overall accuracy of imputation across all sites.

| Coverage | $r^2$ concordance pre-filtering | | $r^2$ concordance post-filtering | |
| --- | --- | --- | --- | --- |
|  | BGI 40X | Illumina 40X | BGI 40X | Illumina 40X |
| BGI 1X | 0.962085 | 0.962852 | 0.970049 | 0.970573 |
| Illumina 1X | 0.961485 | 0.962807 | 0.969826 | 0.970724 |

## Re-use potential

Despite continuous improvements in genotype imputation algorithms, lcWGS imputation remains underutilised as an economical alternative over higher-coverage sequencing. Additionally, the understanding of host genetic markers that predispose to COVID-19 severity is still limited [7].

In this context, our manuscript’s dataset, coupled with the innovative methods we employed, casts a promising light. Not only do we showcase the viability of using lcWGS imputation to generate data for the study of disease-related genetic markers, but also we present a robust validation methodology to ensure accuracy of the data produced. Our ambition is to inspire confidence and stimulate further interest from researchers who wish to deploy a similar approach to a range of other infectious diseases, genetic disorders, or population-based genetic studies, particularly in large scale genomic projects and resource-limited settings where sequencing at higher coverages proves to be prohibitively expensive.

It is important to note, however, that the inherent probabilistic nature of imputed low-coverage genotypes can introduce uncertainty into downstream analyses, and that measures should be taken to mitigate such errors. For example, <u>Petter, Ding</u> [46]Petter, Ding [46] propose a statistical calibration method for polygenic scores (PGS) based on imputed lcWGS genotypes, which is shown to improve estimations of PGS over traditional calculations that ignore genotyping errors in low-coverage sequencing. Understanding the nature of genotyping error is essential to accurately interpret imputed lcWGS results, and, therefore, researchers should be mindful of adopting similar approaches when utilising imputed datasets such as this one.

Beyond the immediate implications in lcWGS imputation, this dataset serves as a valuable resource for investigators studying genetic markers associated with COVID-19 severity. Specifically, the careful methodology we utilised to characterise our patient cohort, through standardised clinical terminology, paves the way for the discovery of genetic components that might be linked to severe COVID-19 disease manifestations and progressions. For instance, this methodology could be tailored to analyse hospitalisation trends in other clinical cohorts, thereby serving as a template for future studies aiming to comprehensively characterise complex diseases.

The analysis of patient hospitalisation, particularly focused on sex and age-related differences, has the potential to inform healthcare policy and clinical guidelines. The insights gained from hospital stay distributions, ICU admissions, and the identification of disease severity across different demographics have broad applications. For example, they could be translated into personalised care strategies or even underpin predictive models that assist healthcare providers delivering more effective treatments [47, 48].

Finally, the validation data regarding comparisons of short-read sequencing platforms is of great importance. As genomic research progresses, the accuracy and reliability of different sequencing platforms becomes increasingly critical. By offering a comparison of imputation accuracy between Illumina and MGI sequencers, we provide an avenue for other researchers to make informed decisions about their sequencing platform. This becomes especially relevant as the scientific community strives to standardise genetic research methodologies [49], to ensure consistent results and comparable outcomes across different studies.

In conclusion, the dataset presented here, though primarily focused on COVID-19 severity, transcends this scope with a broad utility that reaches multiple domains of scientific research. We encourage its reuse hoping that its integration into other studies will advance our collective understanding and response to complex health challenges, such as those presented by COVID-19.

## Availability of source code and requirements

Project name: GLIMPSE low-coverage WGS imputation

Project home page: https://github.com/renatosantos98/GLIMPSE-low-coverage-WGS-imputation

Operating system(s): Linux

Programming language: Bash and Python3

Other requirements: GLIMPSE 1.1.1; samtools 1.16; bcftools 1.16; Python 3.11; numpy 1.24.3; matplotlib 3.7.1; pandas 2.0.3; seaborn 0.12.2; parallel 20230522; mosdepth 0.3.3; plink 1.90b6.21; bc 1.07.1.

License: MIT license

## List of abbreviations

AFR: 1000 Genomes Africans superpopulation
AMR: 1000 Genomes Admixed Americans superpopulation
ARDS: acute respiratory distress syndrome
cM: centimorgans
COVID-19: coronavirus disease 2019
EDTA: ethylenediaminetetraacetic acid
EUR: 1000 Genomes Europeans superpopulation
GP: genotype probability
GWAS: genome-wide association studies
IBD: identity-by-descent
IBS: 1000 Genomes Iberian Populations in Spain population
ICU: intensive care unit
lcWGS: low-coverage whole genome sequencing
MAF: minor allele frequency
PCA: principal component analysis
PGS: polygenic scores
SAS: 1000 Genomes South Asians superpopulation
SARS-CoV-2: severe acute respiratory syndrome coronavirus 2
VCF: variant call format
WGS: whole genome sequencing

## Data Availability

The clinical dataset is available in the European Genome-phenome Archive, under the accession number EGAS00001007573. The other datasets supporting the results of this article are available in our Figshare collection [50] and GitHub repository [51].

## Declarations

### Ethics approval and consent to participate

This study was evaluated and approved by the Clinical Research Ethics Committee of Hospital Clínico San Carlos (code number: 20/313-E_COVID) in Madrid, Spain. In compliance with the provisions of the Declaration of Helsinki and the legislation in force in Spain regarding research with human beings, patients were informed about their participation in this clinical study, clarifying that their participation was voluntary and did not imply any change in their treatment or medical care compared to what they would receive if they did not participate. All patients were informed and voluntary consents were obtained in writing.

### Consent for publication

All data has been anonymised and all links between the identity of patients and the datasets shared in this publication were removed.

### Authors’ contributions

RS and MC conceived the experiment, developed and performed the data processing workflow, performed quality control, and drafted the initial manuscript.

VMT, IP, CdM, OC, VS developed the consent and ethics protocol, recruited the patients, and performed sampling and extraction of purified DNA from samples.

All authors read and approved the final manuscript.

### Authors’ information

N/A

## Acknowledgements

We would like to thank the patients and their families for their cooperation and contribution to this study.

## References

1. Guo G, Ye L, Pan K, Chen Y, Xing D, Yan K, et al. New Insights of Emerging SARS-CoV-2: Epidemiology, Etiology, Clinical Features, Clinical Treatment, and Prevention. Front Cell Dev Biol. 2020;8:410. doi:10.3389/fcell.2020.00410.

2. Tang D, Comish P and Kang R. The hallmarks of COVID-19 disease. PLoS Pathog. 2020;16 5:e1008536. doi:10.1371/journal.ppat.1008536.

3. Severe Covid-19 GWAS Group Genomewide association study of severe Covid-19 with respiratory failure. New England Journal of Medicine. 2020;383 16:1522–34.

4. Covid-Host Genetics Initiative A first update on mapping the human genetic architecture of COVID-19. Nature. 2022;608 7921:E1–E10. doi:10.1038/s41586-022-04826-7.

5. Thibord F, Chan MV, Chen MH and Johnson AD. A year of COVID-19 GWAS results from the GRASP portal reveals potential genetic risk factors. HGG Adv. 2022;3 2:100095. doi:10.1016/j.xhgg.2022.100095.

6. Booth A, Reed AB, Ponzo S, Yassaee A, Aral M, Plans D, et al. Population risk factors for severe disease and mortality in COVID-19: A global systematic review and meta-analysis. PLoS One. 2021;16 3:e0247461. doi:10.1371/journal.pone.0247461.

7. Horowitz JE, Kosmicki JA, Damask A, Sharma D, Roberts GHL, Justice AE, et al. Genome-wide analysis provides genetic evidence that ACE2 influences COVID-19 risk and yields risk scores associated with severe disease. Nat Genet. 2022;54 4:382–92. doi:10.1038/s41588-021-01006-7.

8. Verlouw JAM, Clemens E, de Vries JH, Zolk O, Verkerk A, Am Zehnhoff-Dinnesen A, et al. A comparison of genotyping arrays. Eur J Hum Genet. 2021;29 11:1611–24. doi:10.1038/s41431-021-00917-7.

9. Geibel J, Reimer C, Weigend S, Weigend A, Pook T and Simianer H. How array design creates SNP ascertainment bias. PLoS One. 2021;16 3:e0245178.

10. Brody JA, Morrison AC, Bis JC, O’Connell JR, Brown MR, Huffman JE, et al. Analysis commons, a team approach to discovery in a big-data environment for genetic epidemiology. Nat Genet. 2017;49 11:1560–3. doi:10.1038/ng.3968.

11. Pollie R. Genomic Sequencing Costs Set to Head Down Again. Elsevier, 2023.

12. Meggendorfer M, Jobanputra V, Wrzeszczynski KO, Roepman P, de Bruijn E, Cuppen E, et al. Analytical demands to use whole-genome sequencing in precision oncology. Semin Cancer Biol. 2022;84:16–22. doi:10.1016/j.semcancer.2021.06.009.

13. Alex Buerkle C and Gompert Z. Population genomics based on low coverage sequencing: how low should we go? Mol Ecol. 2013;22 11:3028–35. doi:10.1111/mec.12105.

14. Chat V, Ferguson R, Morales L and Kirchhoff T. Ultra Low-Coverage Whole-Genome Sequencing as an Alternative to Genotyping Arrays in Genome-Wide Association Studies. Front Genet. 2021;12:790445. doi:10.3389/fgene.2021.790445.

15. Gilly A, Southam L, Suveges D, Kuchenbaecker K, Moore R, Melloni GEM, et al. Very low-depth whole-genome sequencing in complex trait association studies. Bioinformatics. 2019;35 15:2555–61. doi:10.1093/bioinformatics/bty1032.

16. Rubinacci S, Ribeiro DM, Hofmeister RJ and Delaneau O. Efficient phasing and imputation of low-coverage sequencing data using large reference panels. Nat Genet. 2021;53 1:120–6. doi:10.1038/s41588-020-00756-0.

17. Nait Saada J, Kalantzis G, Shyr D, Cooper F, Robinson M, Gusev A, et al. Identity-by-descent detection across 487,409 British samples reveals fine scale population structure and ultra-rare variant associations. Nat Commun. 2020;11 1:6130. doi:10.1038/s41467-020-19588-x.

18. Das S, Abecasis GR and Browning BL. Genotype Imputation from Large Reference Panels. Annu Rev Genomics Hum Genet. 2018;19 1:73–96. doi:10.1146/annurev-genom-083117-021602.

19. Browning BL and Yu Z. Simultaneous genotype calling and haplotype phasing improves genotype accuracy and reduces false-positive associations for genome-wide association studies. Am J Hum Genet. 2009;85 6:847–61. doi:10.1016/j.ajhg.2009.11.004.

20. Spiliopoulou A, Colombo M, Orchard P, Agakov F and McKeigue P. GeneImp: Fast Imputation to Large Reference Panels Using Genotype Likelihoods from Ultralow Coverage Sequencing. Genetics. 2017;206 1:91–104. doi:10.1534/genetics.117.200063.

21. Wasik K, Berisa T, Pickrell JK, Li JH, Fraser DJ, King K, et al. Comparing low-pass sequencing and genotyping for trait mapping in pharmacogenetics. BMC Genomics. 2021;22 1:197. doi:10.1186/s12864-021-07508-2.

22. Consortium GP. A global reference for human genetic variation. Nature. 2015;526 7571:68.

23. Lappalainen I, Almeida-King J, Kumanduri V, Senf A, Spalding JD, Ur-Rehman S, et al. The European Genome-phenome Archive of human data consented for biomedical research. Nat Genet. 2015;47 7:692–5. doi:10.1038/ng.3312.

24. Corpas M, de Mendoza C, Moreno-Torres V, Pintos I, Seoane P, Perkins JR, et al. Genetic signature detected in T cell receptors from patients with severe COVID-19. iScience. 2023:107735. doi:10.1016/j.isci.2023.107735.

25. Santos R and Corpas M. Severe COVID-19 Patient Cohort Clinical History. Figshare+. 2023. doi:10.25452/figshare.plus.23932695.v1.

26. Robinson PN and Mundlos S. The human phenotype ontology. Clin Genet. 2010;77 6:525–34. doi:10.1111/j.1399-0004.2010.01436.x.

27. Xiao Y, Zheng X, Song W, Tong F, Mao Y, Liu S, et al. CIDO-COVID-19: An Ontology for COVID-19 Based on CIDO. Annu Int Conf IEEE Eng Med Biol Soc. 2021;2021:2119–22. doi:10.1109/EMBC46164.2021.9629555.

28. Waskom ML. Seaborn: statistical data visualization. Journal of Open Source Software. 2021;6 60:3021.

29. Ewels PA, Peltzer A, Fillinger S, Patel H, Alneberg J, Wilm A, et al. The nf-core framework for community-curated bioinformatics pipelines. Nat Biotechnol. 2020;38 3:276–8. doi:10.1038/s41587-020-0439-x.

30. Garcia M, Juhos S, Larsson M, Olason PI, Martin M, Eisfeldt J, et al. Sarek: A portable workflow for whole-genome sequencing analysis of germline and somatic variants. F1000Res. 2020;9:63. doi:10.12688/f1000research.16665.2.

31. Li H. Aligning sequence reads, clone sequences and assembly contigs with BWA-MEM. arXiv preprint arXiv:13033997. 2013.

32. Chen S, Zhou Y, Chen Y and Gu J. fastp: an ultra-fast all-in-one FASTQ preprocessor. Bioinformatics. 2018;34 17:i884–i90. doi:10.1093/bioinformatics/bty560.

33. Andrews S: FastQC: A Quality Control Tool for High Throughput Sequence Data. http://www.bioinformatics.babraham.ac.uk/projects/fastqc/ (2010). Accessed 16/05/2023 2023.

34. Robbins A. Gawk: Effective AWK Programming. Free Software Foundation Boston; 2004.

35. Van der Auwera GA and O’Connor BD. Genomics in the cloud: using Docker, GATK, and WDL in Terra. O’Reilly Media; 2020.

36. Pedersen BS and Quinlan AR. Mosdepth: quick coverage calculation for genomes and exomes. Bioinformatics. 2018;34 5:867–8. doi:10.1093/bioinformatics/btx699.

37. Van Rossum G and Drake Jr FL. Python reference manual. Centrum voor Wiskunde en Informatica Amsterdam. 1995;40:57–80.

38. Li H. Tabix: fast retrieval of sequence features from generic TAB-delimited files. Bioinformatics. 2011;27 5:718–9. doi:10.1093/bioinformatics/btq671.

39. Danecek P, Bonfield JK, Liddle J, Marshall J, Ohan V, Pollard MO, et al. Twelve years of SAMtools and BCFtools. Gigascience. 2021;10 2:giab008. doi:10.1093/gigascience/giab008.

40. Tange O. GNU Parallel-the command-line power tool. The USENIX Magazine. 2011;36 1:42–7.

41. Belsare S, Levy-Sakin M, Mostovoy Y, Durinck S, Chaudhuri S, Xiao M, et al. Evaluating the quality of the 1000 genomes project data. BMC Genomics. 2019;20 1:620. doi:10.1186/s12864-019-5957-x.

42. Dekeyser T, Genin E and Herzig AF. Opening the Black Box of Imputation Software to Study the Impact of Reference Panel Composition on Performance. Genes (Basel). 2023;14 2:410. doi:10.3390/genes14020410.

43. Sousa da Mota B, Rubinacci S, Cruz Davalos DI, CE GA, Sikora M, Johannsen NN, et al. Imputation of ancient human genomes. Nat Commun. 2023;14 1:3660. doi:10.1038/s41467-023-39202-0.

44. Chang CC, Chow CC, Tellier LC, Vattikuti S, Purcell SM and Lee JJ. Second-generation PLINK: rising to the challenge of larger and richer datasets. Gigascience. 2015;4 1:7. doi:10.1186/s13742-015-0047-8.

45. Hunter JD. Matplotlib: A 2D graphics environment. Computing in science & engineering. 2007;9 03:90–5.

46. Petter E, Ding Y, Hou K, Bhattacharya A, Gusev A, Zaitlen N, et al. Genotype error due to low-coverage sequencing induces uncertainty in polygenic scoring. Am J Hum Genet. 2023;110 8:1319–29. doi:10.1016/j.ajhg.2023.06.015.

47. Molani S, Hernandez PV, Roper RT, Duvvuri VR, Baumgartner AM, Goldman JD, et al. Risk factors for severe COVID-19 differ by age for hospitalized adults. Sci Rep. 2022;12 1:6568. doi:10.1038/s41598-022-10344-3.

48. Wongvibulsin S, Garibaldi BT, Antar AAR, Wen J, Wang MC, Gupta A, et al. Development of Severe COVID-19 Adaptive Risk Predictor (SCARP), a Calculator to Predict Severe Disease or Death in Hospitalized Patients With COVID-19. Ann Intern Med. 2021;174 6:777–85. doi:10.7326/M20-6754.

49. Jeon SA, Park JL, Park S-J, Kim JH, Goh S-H, Han J-Y, et al. Comparison between MGI and Illumina sequencing platforms for whole genome sequencing. Genes & Genomics. 2021;43:713–24.

50. Santos R and Corpas M. Supporting data for “Low-coverage whole genome sequencing for a highly selective cohort of severe COVID-19 patients”. Figshare+ Collection. 2023. doi:10.25452/figshare.plus.c.6347534.

51. Santos R and Corpas M: Low-coverage whole genome sequencing for a highly selective cohort of severe COVID-19 patients. https://github.com/renatosantos98/GLIMPSE-low-coverage-WGS-imputation (2023).

